# Fabrication and Characterization Methods for Investigating Cell-Matrix Interactions in Environments Possessing Spatial Orientation Heterogeneity

**DOI:** 10.1101/2021.05.25.445622

**Authors:** Michael J. Potter, William J. Richardson

## Abstract

Fibrillar collagen is a ubiquitous structural protein that plays a significant role in determining the mechanical properties of various tissues. The constituent collagen architecture can give direct insight into the respective functional role of the tissue due to the strong structure-function relationship that is exhibited. In such tissues, matrix structure can vary across local subregions contributing to mechanical heterogeneity which can be implicated in tissue function or failure. The post-myocardial infarction scar environment is an example of note where mechanically insufficient collagen can result in impaired cardiac function and possibly tissue rupture due to post-MI cellular response and matrix interactions. In order to further develop the understanding of cell-matrix and cell-cell interactions within heterogeneous environments, we developed a method of heterogeneous collagen gel fabrication which produces a region of randomly oriented fibers directly adjacent to an interconnected region of anisotropic alignment. To fully capture and evaluate the degree of alignment and spatial orientation heterogeneity, several image processing and automated analysis methods were employed. Our analysis revealed the successful fabrication of an interconnected spatially heterogeneous collagen gel possessing distinct regions of random or preferential alignment. Additionally, embedded cell populations were observed to recognize and reorient with their underlying and surrounding architectures through our cell-centric analysis techniques.

## 1. Introduction

Fibrillar collagen is a highly abundant matrix component in many tissues and a central determinant of tissue mechanical properties[1]. Fiber orientation and alignment provide a structural basis for mechanical anisotropy, which is essential for the function of many load-bearing tissues, including tendons, ligaments, heart valves, and blood vessels. Additionally, underlying collagen architectures observed in tissues regulate a wide range of varied cellular responses including endothelial cell morphology, migration dynamics in cancer cell populations, and integrin mediated myofibroblast activation[2–5]. In some of these tissues, matrix structure and the resulting mechanical properties vary significantly across local subregions, and the effects of such heterogeneity can be important for tissue function or failure. For example, mechanical stress gradients have been shown to produce heterogeneous structural hallmarks during headfold morphogenesis[6,7]; a hammock-like heterogeneity of matrix orientation from commissure to commissure in heart valves has been a key target of tissue engineering approaches[8,9]; the spatial transition from highly aligned tendon matrix to randomly oriented and mineralized bone matrix at the tendon-bone enthesis is critical to proper load transfer and musculoskeletal function[10,11]; spatial heterogeneity in the structural composition of aneurysms has been associated with high variation in tensile moduli and failure locations[12,13]; and collagen orientation heterogeneity in dermal wounds has been associated with improved cell infiltration and healing rates[14].

Another notable example of heterogeneous collagen orientation is post-myocardial infarction, where failure modes can be related to the mechanical properties of the resultant scar[15]. Myocardial wall rupture post-MI can result from too much degradation (or too little accumulation) of collagen at the infarct location and potentially from misalignment of the fiber architecture[16]. In recent work, we found that stark collagen orientation heterogeneity emerges early post-MI in rat infarcts and remains through at least 6 weeks of the healing time-course[17]. This was true even in cases with low global alignment (i.e., mostly isotropic). More recently, these localized pockets of fiber alignment have predicted increased risk for tissue rupture[18].

In order to circumvent many of the threats presented by mechanically insufficient collagen architecture in scar tissue post-infarction, it is essential to understand how the scar is remodeled by local cell populations. While it is clear that local mechanical stresses play a major role in directing fiber manipulation by local fibroblast populations, many questions remain unanswered regarding cell-matrix interactions in heterogeneous environments such as the following: how much of a role do neighboring fibroblasts have in directing the deposition and fiber manipulation of their neighbors, how does localized regional alignment arise within largely random environments, at what distances can fibroblasts within the collagenous scar environment detect the alignment of the matrix it is embedded within, what effect does the fibroblast have on its neighbors through its own contractile inputs into the matrix network, and many others. As a first step in probing cell-matrix and cell-cell interactions within heterogenous environments, we have developed a method to fabricate heterogeneous collagen gels in vitro with a region of randomly oriented fibers directly adjacent to an interconnected region of anisotropic alignment. In addition, we developed several image processing and automated analysis methods to quantify fiber and cell spatial heterogeneities from fluorescent microscopic live-cell images.

## 2. Methods

### 2.1. Gel Fabrication

Briefly, type I rat tail collagen at a concentration of 4.1mg/ml was purchased commercially (Advanced Biomatrix, San Diego, CA) and mixed on ice with the accompanying kit-provided collagen neutralization solution, 10XMEM, cell-containing media, and Pierce™ streptavidin-coated magnetic microspheres(Thermo Fisher Scientific, Waltham, MA) with an average diameter of 1 μm and at a solution concentration of 10mg/ml prior to pipetting into Nunc™ Lab-Tek™ chambered coverglass slides. First, 41.25 ul of the neutralization solution was pipetted into a microcentrifuge tube followed by 91.8 ul of 10XMEM. For the bead-containing solution, 16 ul of streptavidin-coated magnetic microsphere solution was pipetted into the total solution which was then mixed thoroughly. The addition of the beads is skipped for the bead-less ‘random’ solution. At this point, 371 ul of the rat tail collagen solution was added and mixed with care to avoid microbubble formation. Warm cell-containing media at a volume of 60 ul is then added at a cellular concentration of 16.9 M/ml. Prior to pipetting, dividers were placed within approximately 10.3 mm x 8.3 mm chamber wells to maintain solution separation based on what will become the two regions of interest.

Within each quadrant separated by the dividers illustrated in Fig. 1A, 60 ul of the final mixed solution was pipetted prior to divider removal. The bead-containing ‘aligned’ solution was placed within the outer quadrants while the bead-less ‘random’ solution was pipetted into quadrants located towards the center of the chambered coverglass slide. Aligned solution regions are indicated by exteriorly located blue shading in the chamber slide wells in Fig. 1A. Immediately following pipetting, dividers were removed and the chamber slides containing collagen solution were inserted into a custom-fabricated magnet holder assembly between two 100lb-force neodymium bar magnets (K&J Magnetics, Pipersville, PA) where the gels were polymerized at room temperature for one hour to allow for bead migration and fiber formation. Bead migration along the magnetic field lines imparts the alignment found within the gel as demonstrated in work performed by Guo et al[19]. Following the one-hour room temperature polymerization, gels were placed in standard 37°C, 5% *CO*_2_ incubators.

**Fig. 1.**
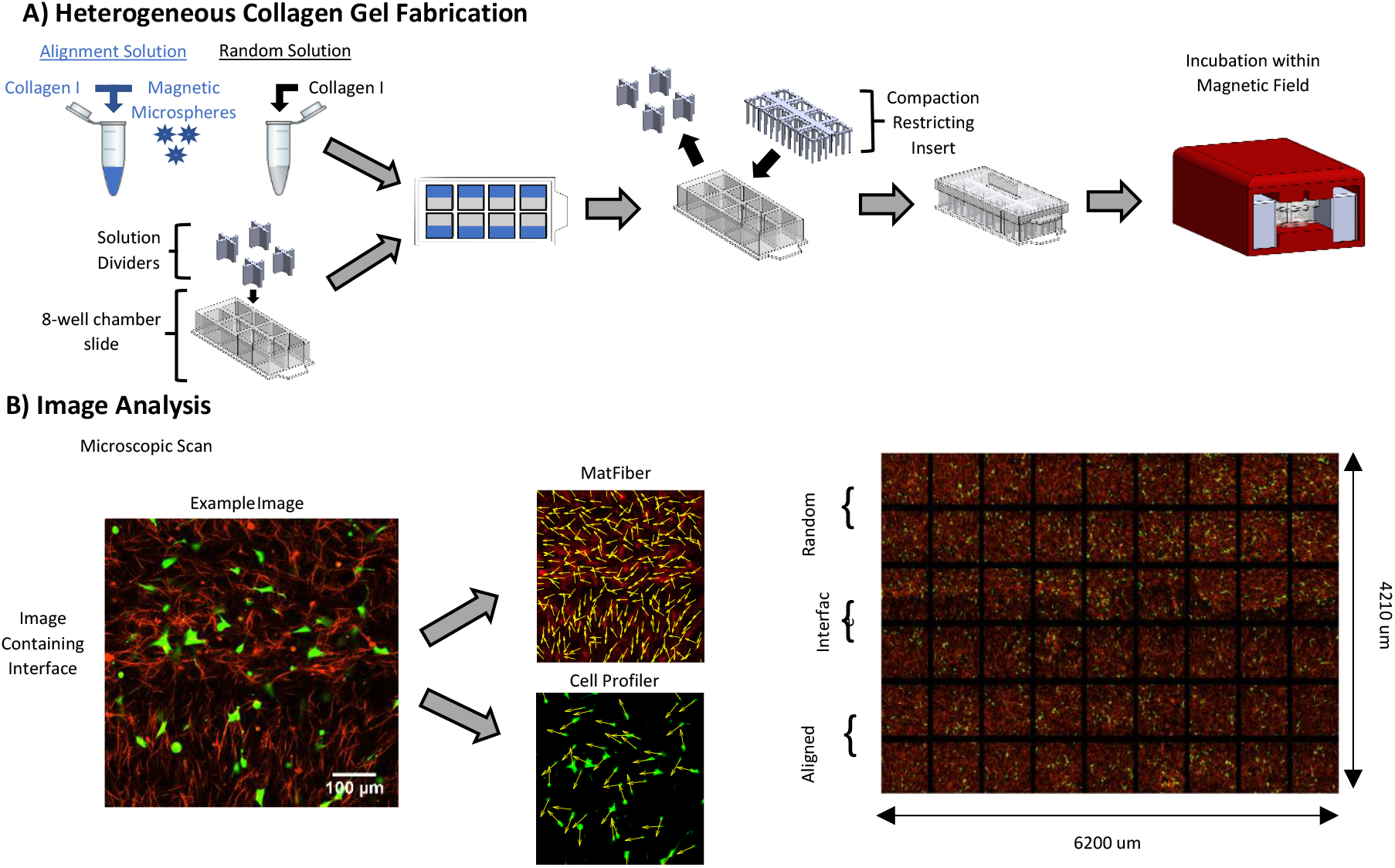
Methodologies Overview: A) Representation of the fabrication process. The two solutions(aligned and random) are prepared and solution dividers are placed within the chamber wells. Aligned solution is extruded into the darker shaded region at the periphery of the well to maintain positioning closer to the external magnets and experience a stronger portion of the magnetic field. Dividers are removed after extrusion, and an insert is placed within the gels to restrict compaction. The assembly is then covered and inserted into the magnet holder assembly. B) An example image is presented to highlight the orientation analyses conducted for fibers and cells within each image. Fiber analysis was conducted on red channel data while cell analysis was conducted on green channel data. The image stitch highlights gel dimensions captured by the 6×9 image grid and exhibits the range of the global coordinates for the gel sample.

GFP-transfected NIH3T3 cells(Cell Biolabs Inc cat: 50-672-593) were cultured in standard culture media (DMEM, 10% FBS, and 1% pen-strep) and suspended within collagen gels with a final collagen concentration of 2.5 mg/ml and a cellular concentration of 1.75 M/ml. One solution contained the addition of streptavidin-coated magnetic microparticles, while the other solution was left without the addition of magnetic microparticles. Gels were maintained with standard culture media which was replaced daily at a volume 1.5x greater than gel solution volume.

Gels were then imaged by confocal reflectance microscopy on days 1, 3, 5, and 7 utilizing a Nikon Eclipse Ti-E motorized inverted microscope with a 20X oil-immersion objective and its accompanying EZ-C1 software(v3.80) for image capture.

### 2.2. Image Acquisition and Processing

Images were taken along a 6×9 grid path that fully captured the center section of each gel sample measuring approximately 6200 μm x 4210 μm. Images were then processed independently using a previously published image processing method(MatFiber [20], code available at http://bme.virginia.edu/holmes) implemented in MATLAB and CellProfiler(v3.1.5) to obtain the orientations (*α*) of fibers and cells within each image. An example image displaying orientation analysis as well as a stitch exhibiting the 6×9 grid path is shown in Fig. 1B. MatFiber quantifies collagen alignment by segmenting the image into a grid of approximately 10 μm x 10 μm subregions, calculating intensity gradients, and returning an angle and image coordinates for each subregion visualized in Fig. 1B via vector field overlay. The implemented CellProfiler pipeline operates by separating foreground from background utilizing a global threshold value. Calculation of the threshold value was performed by the built-in ‘RobustBackground’ thresholding method which assumes an approximate gaussian background distribution and sets the threshold value as the mean plus user determined number of standard deviations of the included pixel range. For our analysis, the included pixel intensity window for threshold calculation ranged from the 20^th^ to the 95^th^ percentile and placed the threshold two standard deviations above the mean of included values. After thresholding, primary objects are identified by size and shape while excluding objects outside of a 12 to 123 micron diameter range. Object coordinates, area, major and minor axis lengths, and orientation are produced. The obtained fiber subregion orientations and coordinate positions as well as the cell coordinate positions, area, major and minor axis lengths, and orientations were placed into tables. Global coordinates were calculated for each cell and fiber subsection based on image and intra-image object position. Calculating global coordinate positions and orientations of cells and fibers enabled us to perform a variety of alignment correlations and regional comparisons across varying spatial ranges and zones as described below.

### 2.3. Fiber Analysis

In order to characterize the fiber architecture transition across the random to aligned regions, we quantified fiber orientation distribution histograms across 10-degree orientation bins from 0 to 180 degrees(n=18 bins). Alignment values were found by calculating the mean vector length(MVL) of fiber angle distributions given by Eq. 1:

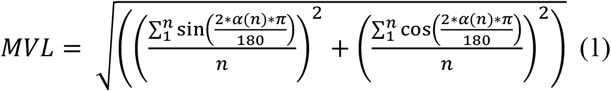

To characterize the spatial distribution of fiber alignment, we grouped fiber angles by their y-position coordinate (i.e., distance relative to the bead-no bead interface) and calculated alignment values for each y-position grouping. To find these averages, the assigned fiber alignment values(MVL) of individual zones were organized by experimental run, treatment, sample, and timepoint before being discretized and binned based on position along gradient. These analysis zones were constructed by segmenting the observed gel section into 210 μm x 210 μm square regions that spanned the coordinate range of data collected. Intra-sample fiber alignment values and zone positions were averaged based on discretized position bins. Alignment value averages were plotted against corresponding gradient position. A two-way ANOVA was conducted for gel type(gradient vs random) and gel position (n = 8). Additionally, in order to evaluate classification of one gel region as ‘aligned’ and the other as ‘random,’ heterogeneous and homogeneous gels were halved, and fiber alignment values were averaged by respective gel region prior to conduction of student t-tests within each gel comparing gel halves.

Evaluation of alignment persistence throughout the time-course was desired to assist in understanding initial local architecture permanence with an embedded cell population. The fiber zones were sorted by experimental run, treatment, sample, and timepoint before discretization and binning by day 1 initial fiber alignment value into eight alignment value ranges each spanning a value range of 0.125. Fiber alignment values within these bins were averaged across samples for each treatment at each timepoint to obtain time-course measurements.

### 2.4. Cell Analysis

In order to characterize the cell recognition of the architecture transition from random to more aligned, we quantified cell orientation distribution histograms across 10-degree orientation bins from 0 to 180 degrees(n=18 bins). We also quantified cell elongation and cell spread area after grouping cells based on the local alignment of collagen fibers. Specifically, cell centric analysis zones were constructed for each individual cell as circular regions with radii length of 210 μm centered on each cell. Cells were grouped based on the local fiber alignment within these zones on day 1, and cell parameters were averaged across these groups for subsequent timepoints. Only cells with a major axis to minor axis ratio of 1.5 were included within analyses utilizing cell orientation to minimize the occurrence of misattributing cellular orientation.

### 2.5. Quantifying Cell and Matrix Heterogeneous Interactions

While our gel fabrication method produced fibrous constructs with clear, macroscale heterogeneity (two starkly different regions), we also sought to test quantification metrics for analyzing more subtle microscale heterogeneities that might exist within each region. To do so, we employed three different approaches. First, we quantified the correlation between cell alignment angles and fiber alignment angles within local pockets of varying sizes. This was accomplished by finding the mean angle (MA) of all cells within a particular range of a central cell, which is calculated by Equation 2:

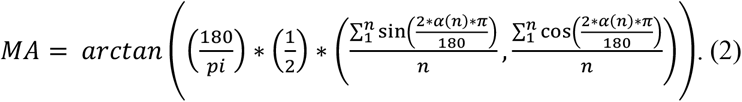

The MA is calculated for all fibers within that same range, then subtracted from the cell MA to quantify a delta mean angle(ΔMA) that represents how closely aligned the cells and fibers are within the region. This was repeated for all cells in each sample, and the MVL of the resulting distribution of ΔMAs was used as an overall measure of localized cell-fiber alignments for a given local zone size. Our second quantification of heterogeneity used a similar approach but analyzed one cell at a time rather than local pockets of cell groups. Specifically, the MA of all fibers within a particular range of a given cell was subtracted from that individual cell’s orientation angle, producing a ΔMA that represents how closely aligned the cell is with the average fiber orientation across a given range. This was repeated for all cells, and the MVL of the resulting distribution of ΔMAs was used as an overall measure of localized cell-fiber alignments for a given local zone size.

Lastly, our third quantification of heterogeneity followed the previous method developed by Richardson and Holmes using alignment vs. distance plots between individual pairs of cells and fibers[18]. Specifically, we calculated the dot product of a cell or fiber’s orientation vector with every other fiber’s orientation vector and then plotted those dot product alignment values against the distance between each pair. We repeated this for thousands of random fiber-fiber and cell-fiber pairings and averaged alignment values within incremental distance bins.

### 2.6. Statistical Methods

Data represent the mean of eight independent replicates and error bars illustrate the standard error of the mean. Statistical analysis performed utilized two-sample t-tests and two-way ANOVAs with n = 8. A p-value was deemed statistically significant when less than 0.05. Frequency plots were constructed by pooling data from the eight independent replicates into a single analysis pool for each respective timepoint and condition.

### 2.7. Code and Data Availability

Analysis code and data freely available at: https://github.com/SysMechBioLab

## 3. Results

### 3.1. Fiber Properties

Due to the importance and prevalence of heterogeneous architecture in collagen-rich environments, we sought to generate continuous tissue constructs possessing intra-gel alignment heterogeneity. In order to evaluate the degree of generated alignment within gel regions at a fundamental level, orientation distributions were constructed with the fiber units of each region of alignment of the heterogeneous gels and illustrated in Fig. 2B. Fibers within the aligned region display a clear peak around the desired 90° orientation bin while the random region exhibits no significant peak. This peak within the aligned region coupled with the lack of distinct peak within the random region again supports the claim that differential alignment within the gel was created. It also appears that the distribution range is maintained through day 7 in both gel sections despite a slight reduction in the peak of the aligned region distribution moving from day 1 to day 3.

**Fig. 2.**
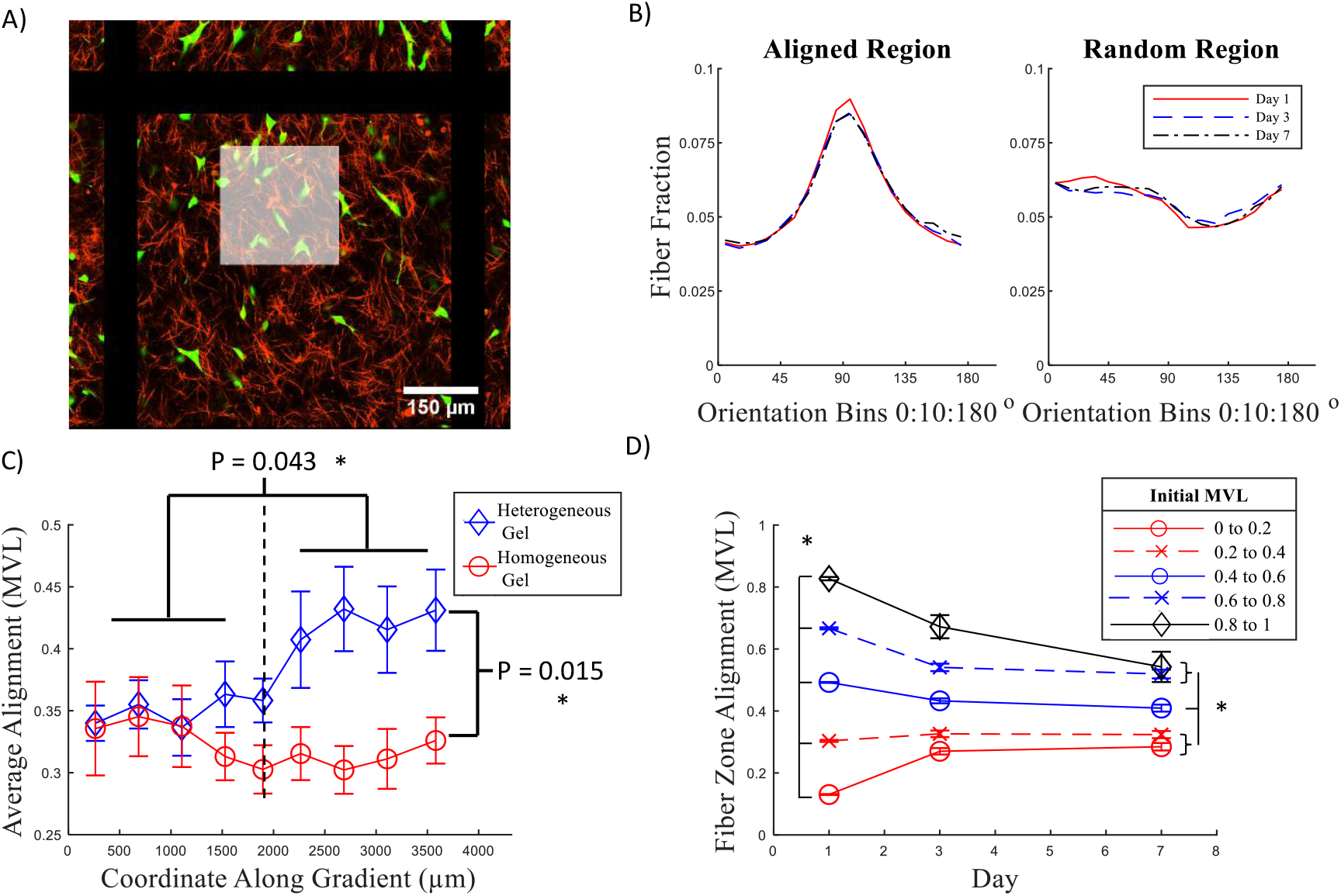
Evaluating Generated Fiber Alignment Heterogeneity. A) A white zone overlay denoting an example zone of analysis that was utilized to evaluate the generation of an alignment gradient. B) Orientation distributions highlighting the differential fiber alignment exhibited by the two interconnected regions of heterogeneous gel constructs. C) Analysis of degree of alignment change across intended gel alignment axis. Statistical values displayed compare aligned region of heterogeneous gel to the random region of the heterogeneous gel and the corresponding homogeneous gel region.(n=8, ±*SE*. *p<0.05). D) Evaluation of fiber architecture maintenance over time-course. Two-way ANOVA revealed significance between all individual day 1 initial alignment groupings while day 7 analysis revealed a loss of significance between the two lowest initial alignment groupings and the two highest initial alignment value groupings. Significance was maintained between the remaining initial alignment groupings.

In order to appropriately assess the generation of a heterogeneous architecture, we next evaluated the average subregion degree of alignment across the gel along the intended gradient. Instead of evaluating hand selected pockets from each greater region, observing alignment across the entirety of the gel enabled a clearer picture of whether the desired spatial trend was generated. An increase in degree of alignment is discernible from the alignment evaluation illustrated in Fig. 2C. Progressing across the gel reveals a clear deviation away from the alignment values exhibited by analysis of purely random control gels while the random region of the gel possesses alignment values within the range of a purely random gel. A two-way ANOVA was conducted comparing discretized cross-gradient alignment. The analysis revealed no difference based on which gel, heterogeneous or homogeneous, was analyzed. This is likely due to the gradient gel possessing a region of random orientation as well as a region of greater alignment. This translates into recognizing no real difference between samples at the full gel level. However, there was significance at the p = 0.01 level when considering position within the gel along the axis of the gradient, suggesting a confirmation of the generation of a region of more aligned fibers. Additionally, to determine if there was not only a gradient generated, but also sufficient alignment to categorize gel halves as separate alignment regions, gel halves were categorized as either a ‘Random’ or ‘Aligned’ region. Then 210um analysis zones within each region were averaged to obtain an overall value for the respective gel halves. T-tests were then performed to compare the regions and the resulting p-values are displayed in Fig. 2C. Significance at the p = 0.05 level was observed when comparing the two regions of the heterogeneous gel as well as when comparing the aligned region of the gradient gel to both regions of the random gel. No significance was observed between the random region of the gradient gel and either region of the random gel. This supports the evidence that a gradient was generated across the heterogeneous gels through the streptavidin magnetic microparticle migration method.

In addition to evaluating whether the desired architecture was generated, maintenance of architecture was an additional topic of interest as embedded cell populations can manipulate their environments when they engage with them[21]. Fiber subregions were grouped by initial alignment value and average alignment was found for each grouping. These groupings were maintained for analysis of day 3 and day 7 degree of alignment as well. Resultant values are shown in Fig. 2C and exhibit a trend toward centrality of alignment values. The greatest changes in average alignment are observed day 1 to day 3 while day 3 to day 7 is relatively stable. Two-way ANOVA revealed statistical significance between all day 1 values while day 7 analysis revealed a loss of significance between the two lowest initial alignment groupings and between the two greatest alignment groupings while significance is maintained concerning comparison to the other initial alignment value groupings. We attribute the change in alignment values to actions of the embedded fibroblast populations onto their surrounding environment. The trend toward centrality may suggest cellular detection of surrounding architecture through imparting tension onto and detecting tension on the matrix.

### 3.2. Cell Response to regional architecture

In addition to characterizing the architecture of the gel, cell response to architecture is also of interest as cell morphologies are directed by their environments[22–25]. The distribution of cellular orientation was generated by region for each day of observation in Fig. 3B to highlight any trends in cellular orientation related to gel region. A peak is observed throughout the experimental time-course within the aligned region which is maintained through day 7. Within the random region, the distribution appears less consistent through the range of angles and does not exhibit a clear orientation preference. These orientation distributions resemble the underlying fiber orientation distributions shown in Fig. 2B for each respective region of the gel.

**Fig. 3.**
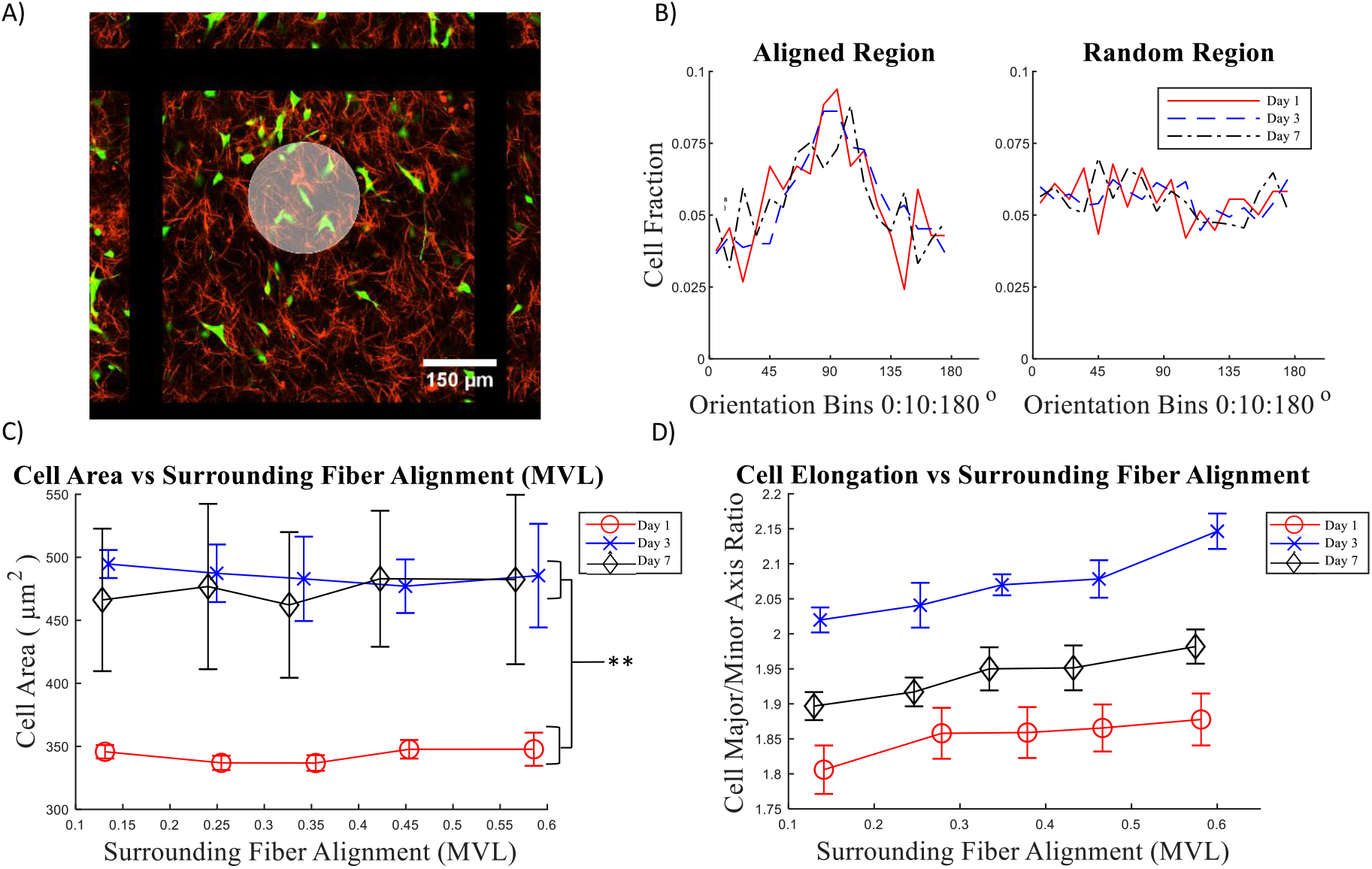
Demonstrating Cell-Centric Analysis Through Orientation and Parameter Response to Matrix Environment. A) A white zone overlay denoting an example zone of analysis that was utilized to support cell-centric analysis methods. B) Orientation distributions highlighting the differential cellular alignment exhibited within the two interconnected regions of heterogeneous gel constructs. C) Analysis of cell area by day compared to the level of alignment exhibited by local collagen fiber architecture within heterogeneous gel constructs(n=8, *±SE.* *p<0.05, **p<0.01). D) Analysis of cell elongation by day compared to the level of alignment exhibited by local collagen fiber architecture within heterogeneous gel constructs.

Fig. 3C highlights the difference in cell area between day 1 and the remainder of the experiment. Low cell area at day 1 suggests that cells appear to have not spread and begun interacting with their environment until some point between the initial day 1 and following day 3 observations as day 1 spreading values across all quintiles exhibited significant difference at the p = 0.01 level compared to all day 3 and day 7 values. Full spreading appears to have occurred by day 3 and maintained through day 7.

Quantifying cell elongation in Fig. 3D indicated a significant increase in elongation from day 1 to day 3 and day 7. While a slight positive relationship appeared between elongation and local collagen fiber alignment for days 1, 3, and 7, no intra-day significant difference was detected by two-way ANOVA.

### 3.3. Fiber-Cell Interactions

A central motivation for our work was to assess the distance at which embedded cell populations can detect and direct regional alignment and coordinate with neighboring cells. Fig. 4B highlights a clustering of delta values around 0 showing a general agreement amongst cell and fiber mean angles between zones. Concerning the smaller analysis zone sizes, over 43% of the orientation deltas fall between −10° and 10°.This agreement is seemingly enhanced in the larger zone size as a greater frequency is noticeable around the smaller delta value bins. Greater than 67% of the orientation deltas for the larger zone analysis falls between −10° and 10°. This means a higher portion of analysis zones have small differences between fiber and cell mean angles when considering large analysis zones. When applying this analysis to the central cell upon which the zones are anchored and finding the delta between the central cell’s orientation and the mean angle of the surrounding fibers, the resulting distribution once again peaked at the lower delta values corresponding to rough agreement of orientation as seen in Fig. 4C. However, increasing the zone size has a marginal effect on agreement between cell orientation and fiber zone orientation. The proportion of deltas falling between −10° and 10° for the smaller and larger zone sizes respectively are 31.5% and 26.6% meaning the opposite trend occurred when comparing to the delta distributions for mean angles of cell and fiber populations within the zones.

**Fig. 4.**
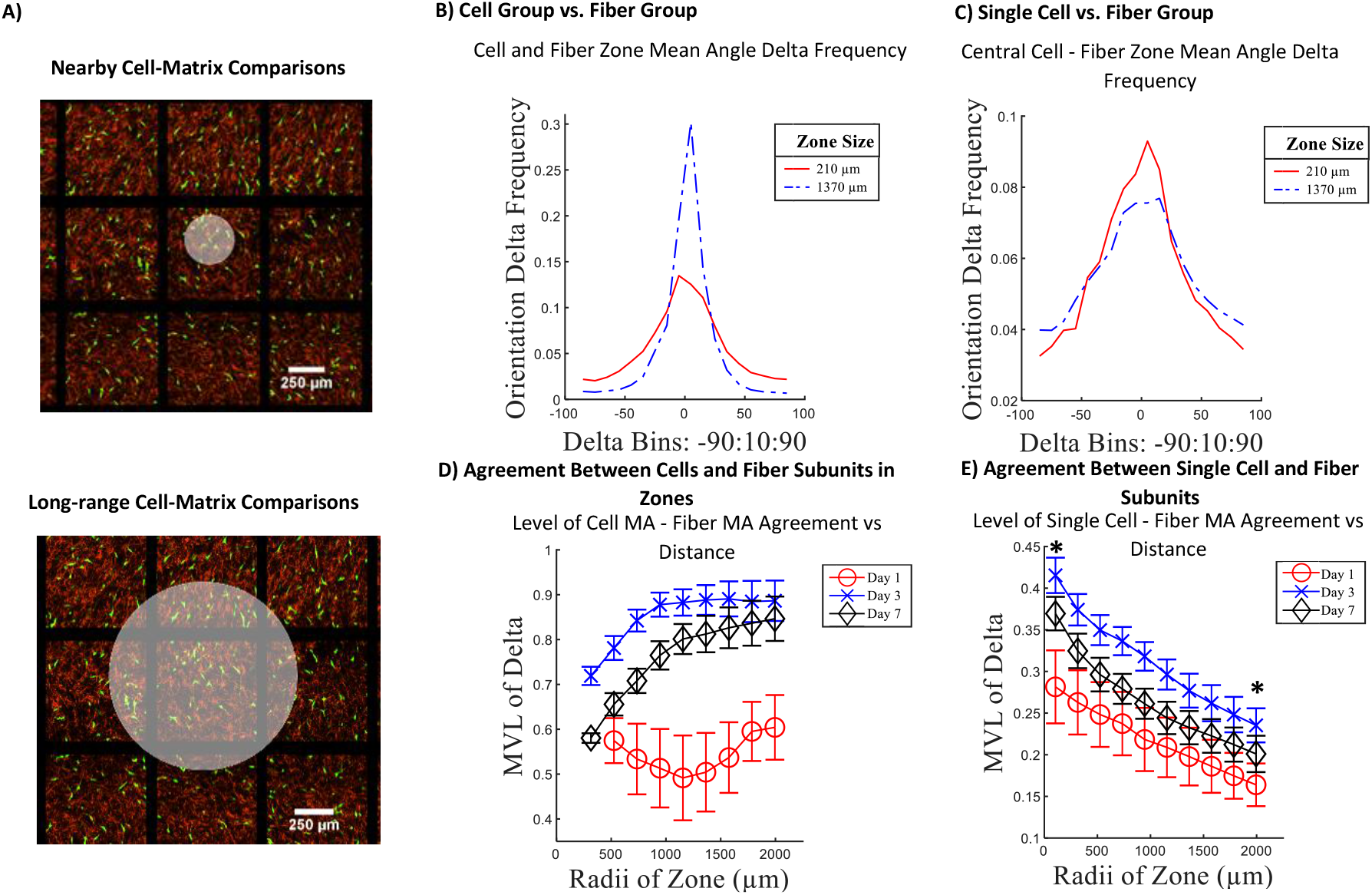
Determining Cell-Matrix Interactions and Recognition Across Varied Distances. A) Example white overlay zones highlighting varied cell-centric analysis zone sizes. B) Analysis of mean angle orientation delta frequency of cells and fibers included in zone between two distinct zone sizes. C) Central cell orientation and surrounding fiber mean angle delta frequency for two distinct zone sizes. D) Evaluating the uniformity of orientation agreement of cells and fibers included in zone in relation to size of analysis zone. E) Evaluating the uniformity of orientation agreement of central cell and surrounding fiber mean angle in relation to size of analysis zone. Significance was observed when comparing the smallest and largest zone sizes of the analysis for day 3 and day 7(n = 8, ±*SE*. *p<0.05).

In addition to analyzing distributions, calculating the mean vector length of the delta distributions of increasing zone sizes provides better insight into the uniformity of the orientation deltas as you increase the zone size. As zone size is increased, there is more likely to be a uniform deviation between cell zone mean angle and fiber zone mean angle. This is observed in the gradual positive slope exhibited by the day 3 and day 7 plots in Fig. 4D. A two-way ANOVA revealed a significant difference between days analyzed, but no significance for radii of zone within each day. For single cell analysis, as zone size is expanded, the agreement of delta values found decreases. With larger zone sizes, there is less likely to be a uniform deviation between central cell orientation and surrounding fiber orientation. Multiple comparisons ad-hoc analysis following two-way ANOVA evaluating the dataset by distance and day did reveal a significant difference between MVL values found for the smallest zone size and the largest zone size for both days 3 and 7 with p-values less than 0.05. To further evaluate the generation of heterogeneity within our collagenous gel constructs and the cellular recognition of this architecture, individual fiber subunits and cells were compared to other fibers within a certain distance by computing the dot product of the central analysis object’s(cell or fiber) orientation with each individual zone fiber’s orientation within the analysis range. These dot products were averaged and plotted against the respective distance revealing a greater level of orientation agreement for close-range objects and a decline in agreement as distance was increased. Cell agreement with individual fibers is shown in Fig. 5A while fiber to fiber agreement is shown in Fig. 5D. Two-way ANOVA revealed significance by distance and day for both cell to fiber and fiber to fiber analyses with p-values drastically lower than p = 0.01. This particular analysis also enables an approximation for how far cells and fibers are locally aligned as all curves tend to flatten around 3000 μm.

**Fig. 5.**
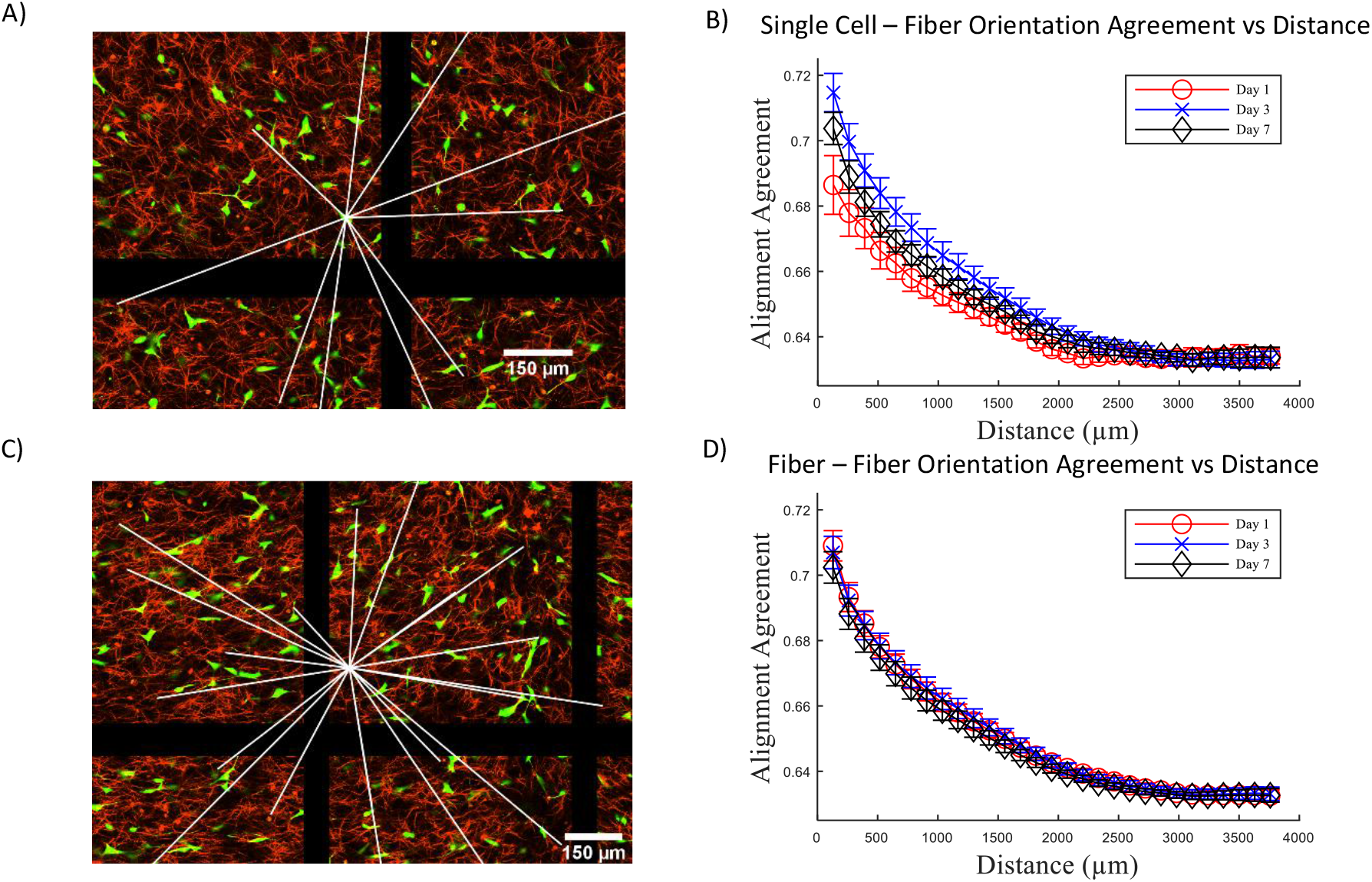
Object to Object Anisotropic Orientation Agreement Decreases with Distance. A,C) Example image with white line overlay illustration denoting object to object analysis; cell to fiber and fiber to fiber respectively. B) Degree of alignment agreement calculated via dot product between single cells and fibers in relation to distance between analysis objects. D) Degree of alignment agreement calculated via dot product between single fiber and other fibers within construct in relation to distance between analysis objects.

## 4. Discussion

In this work, we present a simple methodology to generate collagen gel constructs possessing neighboring regions of isotropic and anisotropic fiber orientations. This platform can be utilized to probe spatial and temporal interactions of cells with fibrous matrix architectures in vitro that could inform further experimentation aimed at understanding cell behavior across neighboring regions possessing inherent differences in architecture. Through our zone-based analysis methodology, it was determined that not only did the gels possess a clear gradient of alignment across the desired dimension, but the angular distribution of fibers within each region was representative of the desired preferred alignment. Over the course of a week, subregion orientation analysis revealed a general trend toward centrality throughout the construct with the largest changes occurring between day 1 and day 3 as the embedded fibroblast populations spread, engaged with, and acted upon their surrounding environment. This process is easily discernible within the analysis of cell orientation, elongation, and spreading. While very little spreading and elongation was observed on day 1, analysis revealed a discernible orientation preference coinciding with the orientation of bead migration and the resultant underlying fibrous architecture. While a recognition of their surrounding environment is evident from a simplified cell parameter analysis, directly analyzing the degree of agreement between a central cell and fibers contained within varied analysis zone sizes revealed a rapid decrease in relevancy for mean fiber angle and distance from the central cell. However, when evaluating the mean orientation of the population of cells within the zone against the mean orientation of fibers, the agreement was improved with increasing zone size. This highlights the complicated nature of translating microscopic relationships to macroscopic behaviors and the importance of assessing matrix organization at the particular scales most relevant to a specific question of interest.

Combining heterogeneity fabrication techniques with heterogeneity analysis metrics could enable many future studies evaluating the role fibroblasts play in directing neighboring cells through time-lapse analysis of cell morphology and various molecular markers compared to neighboring cell expression of target proteins of interest. Analysis from a matrix-focused perspective can lead to insights regarding the architecture of the matrix by closely analyzing the time-course of local pocket degree of alignment arising within aligned or random regions as a result of embedded cell population stimulation and underlying fiber architecture provides this type of architecture minded analysis. These concepts of neighbor-informed and regional remodeling are partly inspired by our previously conducted computational modeling studies analyzing the spontaneous emergence of fiber orientation heterogeneity through long-, medium-, and short-range sensing of fiber orientations by embedded cell populations. In these computational analyses, long-range sensing did result in cell and fiber orientation heterogeneity resembling experimental infarct data[18]. Exploring these computational results through experimental means utilizing our simple fabrication platform can enable the elucidation of the underlying mechanisms of the rise of heterogeneous fibrous architecture in relevant systems.

While other alignment methodologies exist for producing aligned constructs, such as microfluidic fabrication, drop casting, electrospinning, and bioprinting, not all of them can produce a continuous heterogeneous construct, and the methodologies that can are typically complicated, intricate, time consuming, or at the least produce a limited range of smaller construct sizes. Microfluidic fabrication of aligned collagen matrices typically involve additional polymer inclusions and result in very aligned constructs that lack architecture heterogeneity which are useful for the in vitro simulation of highly aligned tissues but lack applicability towards highly heterogeneous tissues.[26;27] Electrospinning methods have recently been shown capable of generating gradients of alignment through fibrous constructs.[28] This methodology is however limited to certain compatible polymer solutions limiting overall application to experimental methods desiring heterogeneous collagen gels possessing no additional polymer inclusions. Drop casting was demonstrated to generate collagen gels possessing varying degrees of alignment throughout the gel constructs in the radial direction from the placed collagen droplet.[29] This methodology appears to be useful for the generation of many replicates of like-collagen gels, but the radial nature of the alignment could confound and limit analysis methods intended to elucidate cooperative cellular response and the fabrication requires intricate environmental control. Bioprinting methodologies, like other methodologies mentioned, are capable of generating constructs with controlled anisotropy and are very promising for intricate design of larger and complicated constructs.[30] However, bioprinting methodologies require specialized fabrication tools possessing inherently intricate levels of control. The methodology we have presented here simply requires typical collagen gel preparation, magnetic microparticle inclusion, a support structure, and a brief room temperature delay within a magnetic field prior to incubation. Fabrication yields intra-gel alignment heterogeneity of large volumetric constructs.

A limitation to this methodology is the lack of intricate spatial control. While a transition threshold is generated between isotropic to anisotropic architecture, more elaborate and smaller scale heterogeneities were not fabricated. An additional limitation is the uppermost levels of alignment achieved. While certain smaller regions possessed alignment levels above the 0.8 level on the scale of 0 to 1, bulk alignment of the aggregate were much more moderate. If extreme anisotropy in architecture is desired, then it may be more appropriate to utilize one of the other previously mentioned alignment methodologies. It may be possible to generate more complex patterns of alignment within fabricated collagen gels through the employment of electromagnets and more complex rigs that adjust the position, strength, and direction of the magnetic field in relation to the fabricated gels. There are many more possible experimental variations to pursue through the utilization of easily fabricated heterogeneous collagen constructs paired with our various analysis considerations.

## 5. Conclusion

Spatial orientation heterogeneity of collagen plays an important role in the relationship between micro-architecture and macro-functionality of tissues. While many methodologies capable of generating anisotropic alignment exist, there are trade-offs concerning time, cost, complexity, spatial control, and purity of included matrix components. This work presents a simple and effective heterogeneous collagen gel fabrication and automated analysis methodology. Spatial orientation heterogeneity was verified through regional evaluation of fiber orientation and cross-construct degree of alignment. Additionally, cell-centric methodologies employed highlight the importance of evaluating cellular response in the context of not only its immediate surrounding environment but also in the context of its environment at greater distances. This platform enables efficient and low-cost experimentation probing fundamental cell-matrix interactions within spatially heterogeneous environments.

## Acknowledgements

We would like to thank our fellow members of the Systems Mechanobiology Lab for useful discussions and support. We also thank the Clemson Light Imaging Facility for the availability of and assistance with their confocal reflectance microscopy instrumentation.

